# A unique intestinal epithelial cell that expresses canonical lymphoid genes

**DOI:** 10.64898/2026.10.21.760658

**Authors:** Swarnabh Bhattacharya, Krithika Badarinath, Guodong Tie, Pratik N. P. Singh, Onur Eskiocak, Ruiyang He, Bin Bao, Su-Hyung Lee, Scott Snapper, Kun Huang, James R. Goldenring, Semir Beyaz, Ramesh A. Shivdasani

## Abstract

Enterocytes and intestinal secretory cells harbor innate immunity. However, certain transcriptional programs –even beyond rearranged T- and B-cell receptor genes– are considered the exclusive purview of adaptive immune cells, including abundant and diverse intestinal intra-epithelial lymphocytes (IELs). Here, we identify and characterize columnar intestinal secretory cells that co-express classic epithelial and lymphoid genes, including *Ptprc*, *Cd3e*, *Cd8a*, and *Gzma/b* but not T- or B-cell receptors. These previously unrecognized epithelium-derived lymphoid-like cells (EDLCs) differ from any known resident cell type, including IELs or tuft cells. Rare in healthy mouse and human intestines, their numbers increase upon IL-25 stimulation or IL-10 receptor deficiency *in vivo* and upon IL-13 stimulation *in vitro*. EDLC chromatin is uniquely accessible at thousands of *cis*-elements, including lymphoid and epithelial loci. These findings reveal a transcriptionally and epigenetically distinct intestinal epithelial derivative that expresses prominent lymphoid character.

## Introduction

Self-renewing Lgr5^+^ intestinal stem cells (ISCs) in crypt bottoms^1^ continuously generate absorptive enterocytes and fewer secretory cells.^2^ Although intestinal epithelial cells are known to display innate immunity^3,4^ and present foreign antigens,^5^ none are known to express markers classically associated with adaptive T-cell immunity, such as CD3, CD4, CD8, and Granzymes. Intestinal expression of these genes is restricted to *Ptprc^+^* (CD45^+^) intra-epithelial lymphocytes (IELs), which originate in the thymus or bone marrow,^6,7^ cross the lamina propria, intercalate between epithelial cell bases, and constitute at least 10% of the epithelium.^8–10^ “Natural” IELs (TCRαβ^+^CD8αα, TCRyδ^+^CD8αα, and TCR^−^innate lymphoid-like cells (ILCs)) home to the intestine soon after arising in the periphery, while “induced” IELs (conventional CD4^+^ or TCRαβ^+^CD8αβ^+^) first encounter antigens elsewhere.^7,11,12^ Here we report the existence of previously unrecognized ISC-derived mouse columnar secretory cells that express both epithelial (e.g., *Epcam*, Keratins) and T lymphocyte (e.g., *Ptprc*, *Cd3e*, *Cd8a, Gzma, Gzmb*) genes but not T- or B-cell receptors. These rare EDLCs are transcriptionally and epigenetically distinct from other intestinal cell types, including tuft-2 cells or IELs, and they do not represent phagocytosis, efferocytosis, or lymphocyte-epithelial fusion events. Rather, they constitute a distinct secretory cell type whose numbers increase in conditions that trigger a type 2 immune response (IL-25 *in vivo*, IL-13 *in vitro*)^13^ and under the inflammatory state present in Il10r^−/−^ mice.^14,15^ Human intestines contain similar cells.

## Results

### Identification of intestinal epithelial cells with a unique dual phenotype

The intestinal secretory lineage diversifies by restricting certain transcription factors (TFs) to selected daughters of Lgr5^+^ ISCs. *Atoh1* and *Neurog3*, for example, are confined to goblet/Paneth or enteroendocrine (EE) cells and required for their respective genesis^16,17^ (Fig. 1A). As these TF genes express early in secretory differentiation, the cells labeled when Cre recombinase is expressed under their control are dominated by the cognate cell type but include other secretory cells and even some ISCs;^18,19^ however, lack of *Atoh1* or *Neurog3* in mesenchymal or lymphoid cells precludes labeling of non-epithelial cells. In previous characterization of the secretory lineage, labeling with *Atoh1^Cre^*(Ref. ^18^, Fig. S1A) or *Neurog3-Cre* (Ref. ^20^, Fig. S1B) consistently identified small single-cell (sc) populations that clustered apart from others. In both cases, genes that distinguish this unique population, e.g., *Cd3e*, *Cd8a*, *Gzma/b*, and *Ptprc* (cell surface marker *Cd45*), are classic lymphocyte markers absent from epithelial cells (Fig. 1B). Upon merging the *Neurog3^Cre^*-labeled population into the *Atoh1^Cre^* dataset, the unusual cells clustered together (Fig. 1C) and based on their identification by epithelial lineage tracing and expression of lymphoid genes, we call them epithelium-derived lymphoid-like cells (EDLCs). Although both FACS isolation and scRNA microfluidics exclude cell clumps, we also used the scDblFinder package for computational exclusion of possible heterotopic doublets,^21^ which were not enriched among EDLCs (Fig. 1C and Fig. S1C). Moreover, doublets or fused cells that evaded these filters would be expected to carry unique RNAs from two distinct cell types, but neither *Atoh1*- nor *Neurog3*-labeled EDLCs had an excess of unique transcripts (Fig. 1C). In addition to classic lymphoid markers and lymphoid-specific TFs (e.g., *Runx3* and *Ikzf2*), EDLCs expressed epithelium-restricted genes (e.g., *Epcam*, *Krt8*, *Krt18*) and secretory markers such as *Muc2*, *Fcgbp*, *Lyz1* and *Defa24* (Fig. 1D).

**Figure 1.**
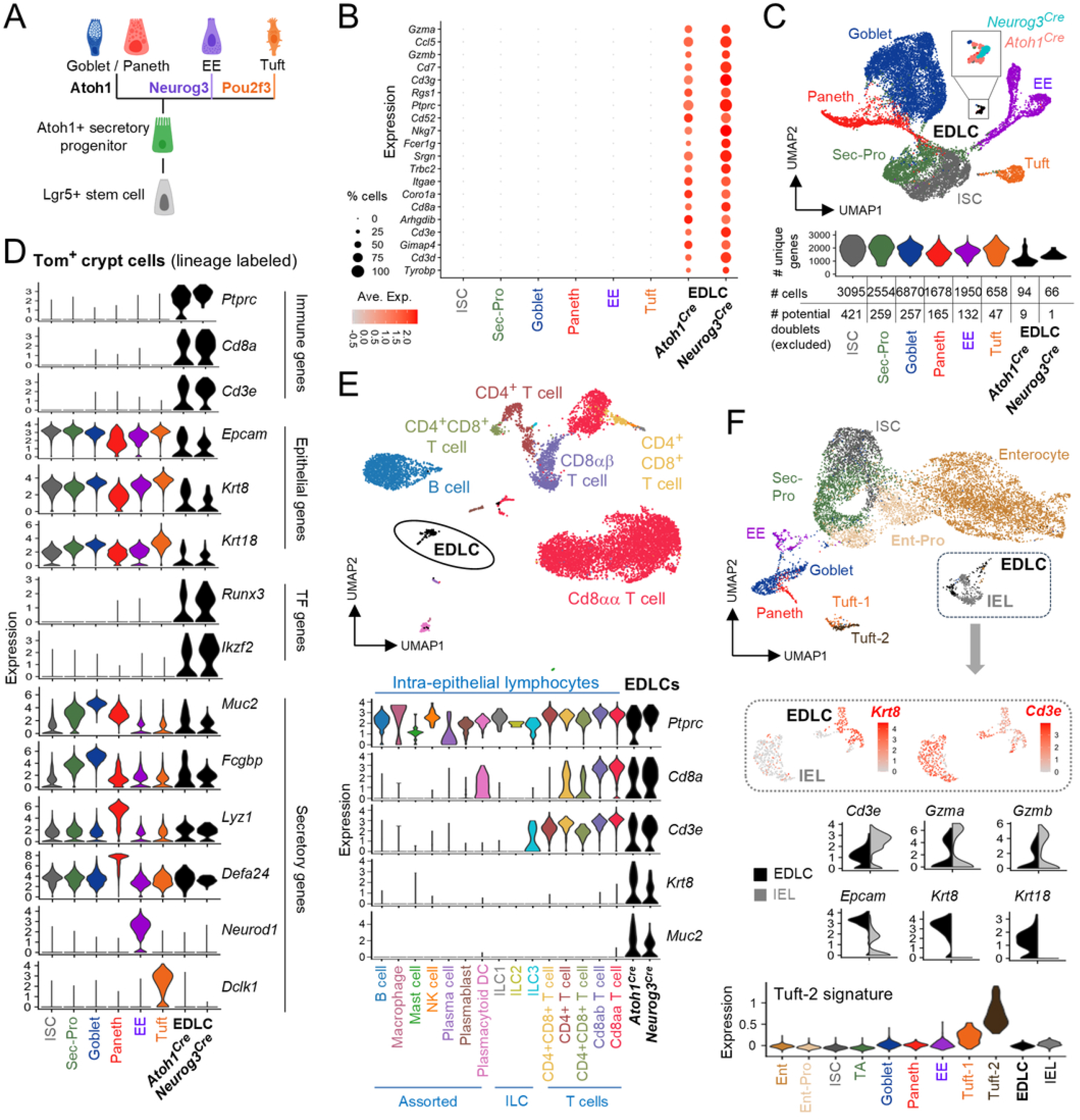
Identification and characterization of epithelium-derived lymphoid-like cells (EDLCs) by scRNA-seq. **(A)** Schema of intestinal epithelial secretory differentiation. **(B)** Top 20 genes enriched in distinct, *Atoh1*-labeled or *Neurog3*-labeled clusters of epithelium-derived lymphoid like cells (EDLCs, Figs. S1A and S1B) reflect a T-cell program. Circle diameters: within-cluster probability of gene detection, fill shades: normalized mean expression level. ISC, intestinal stem cell; Sec-Pro, secretory progenitor; EE, enteroendocrine. **(C)** UMAP plot of *Atoh1*-labeled crypt cells merged with *Neurog3*-labeled EDLCs. EDLCs from both sources combined are shown in black and magnified in the box to show *Atoh1*-derived (salmon) and *Neurog3*-derived (teal) cells. The violin plot displays numbers of unique genes identified in each scRNA-seq cell cluster. EDLCs express fewer genes (reflecting sequencing of fewer cells) and lack the elevated numbers expected in cell doublets. The table displays numbers of cells and potential doublets (predicted by scDblFinder and removed from all analyses, including Figs. 1B and 1D), indicating that EDLCs are not a spurious doublet artifact. **(D)** Pseudobulk-averaged expression of lymphoid (*Ptprc*, *Cd8a*, *Cd3e*), epithelial (*Epcam*, *Krt8*, *Krt18*), transcription factor (*Ikzf2*, *Runx3*) and secretory (*Muc2*, *Fcgbp*, *Lyz1*, *Defa24*, *Neurod1*, *Dckl1*) genes, revealing the dual character of EDLCs. **(E)** Merged UMAP plot of intestinal intraepithelial immune cells ^24^ and *Atoh1*-labeled ^18^ and *Neurog3*-labeled ^20^ EDLCs. Below, pseudobulk-averaged expression of lymphoid (*Ptprc*, *Cd8a*, *Cd3e*), epithelial (*Krt8*), and intestinal secretory (*Muc2*) genes reveals lack of epithelial markers in IEL cells and expression of lymphoid and secretory markers in EDLCs. NK, natural killer; ILC, innate lymphoid cells. **(F)** UMAP plots from scRNA analysis of whole intestinal epithelium (GEO series GSE210724; all potential doublets removed, Fig. S2B). Isolation of IELs and EDLCs from the totality of cells shows that they fall in distinct sub-clusters. Representative features demonstrate lymphoid gene (*Cd3e*, *Gzma*, *Gzmb*) expression in both IELs and EDLCs but epithelial gene (*Epcam*, *Krt8*, *Krt18*) enrichment only in EDLCs. Below, pseudobulk-averaged expression of the tuft-2 signature (top 100 differential genes) ^22^ indicates that EDLCs lack these transcripts, distinguishing them from tuft-2 cells.

*Cd3e*, *Cd8a*, *Gzma/b*, and other genes mark T lymphocytes, but *Ptprc* is a broad blood-cell marker reportedly also expressed in tuft-2, a tuft cell subpopulation.^22^ Using published markers^22^ to distinguish tuft-1 from tuft-2 cells in *Atoh1*-labeled crypts, we found that EDLCs lacked a tuft-2 cell signature or specific tuft markers and, conversely, that tuft-2 cells lacked lymphoid genes (Fig. S1D). Nor did EDLCs match the profile of rare epithelial microfold (M) cells^22,23^ (Fig. S1E). Moreover, *Ptprc* expression was restricted to EDLCs and negligible in 611 tuft cells (Fig. 1D and Fig. S1D) and *in vivo* immunostaining revealed PTPRC in abundant lamina propria cells but not in epithelial cells expressing the canonical tuft marker DCLK1 (Fig. S2A). Thus, EDLCs differ from any known intestinal epithelial cell; marker genes suggest greater similarity to goblet/Paneth than to EE or tuft cells (Fig. 1D).

To distinguish EDLCs from IELs, we normalized data from *Atoh1*-labeled and *Neurog3*-labeled EDLCs together with scRNA-seq data from a published study of all IEL types.^24^ Within the resulting plot of transcriptional similarities among cells, EDLCs from both mouse strains combined into a distinct *k*-nearest neighbor (KNN) cluster (Fig. 1E). While sharing expression of classic lymphoid genes with IELs, especially T cells, EDLCs also expressed distinct epithelial and secretory markers absent from any IEL population (Fig. 1E). Conversely, we examined scRNA-seq data from wild-type mouse villus and crypt epithelium combined (GEO series GSE210724), i.e., independent of *Atoh1* or *Neurog3* labeling, again excluding potential doublets using scDblFinder^21^ and distinguishing tuft-1 from tuft-2 cells (Fig. 1F and Fig. S2B). EDLCs clustered near IELs but were readily distinguished from the latter by dual expression of lymphoid and epithelial genes and they lacked the tuft-2 mRNA signature (Fig. 1F). EDLC transcripts included epithelial TFs (Fig. S2C). A paucity of secretory cell products in villus-predominant EDLCs contrasted with their abundance in crypt cells (compare Fig. 1D and Fig. S2C), implying that villus EDLCs diminish their secretory vestiges.

### EDLC detection and characterization independent of scRNA-seq

To identify EDLCs independent of scRNA-seq and knowing that the secretory lineage derives from Lgr5^+^ ISCs (Fig. 1A), we administered tamoxifen to mice that carry the *R26R^Tom^* reporter gene^25^ and different Cre-drivers, followed 1.5 days (*Atoh1^Cre^*) or 5 days (*Lgr5^Cre^*, Ref. ^1^) later by crypt cell flow cytometry. In each case, a distinct CD45^+^ cell population represented ∼0.5% of tdTom^+^ cells and 0.05% to 0.08% of all epithelial cells, and CD45^+^ or CD8^+^ IELs (tdTom^−^) outnumbered these EDLCs (Fig. 2A). At least two-thirds of EDLCs, distinguished from IELs by EPCAM expression, also expressed the surface marker CD8A (Fig. 2A), consistent with *Cd8a* mRNA expression in 50-75% of individual EDLCs (Figs. 1B and 1D). By light scatter characteristics and microscopic examination of FACS-sorted cells, EDLCs (tdTom^+^ CD45^+^) were larger than IELs (tdTom^−^ CD45^+^) and similar in size to other *Atoh1^Cre^*-labeled CD45^−^secretory cells (Fig. 2B). EDLCs carried single nuclei, similar in size to those of epithelial cells and larger than those of IELs (Fig. 2B), further indicating that they do not represent fused cells, which are often binucleate.^26,27^

**Figure 2.**
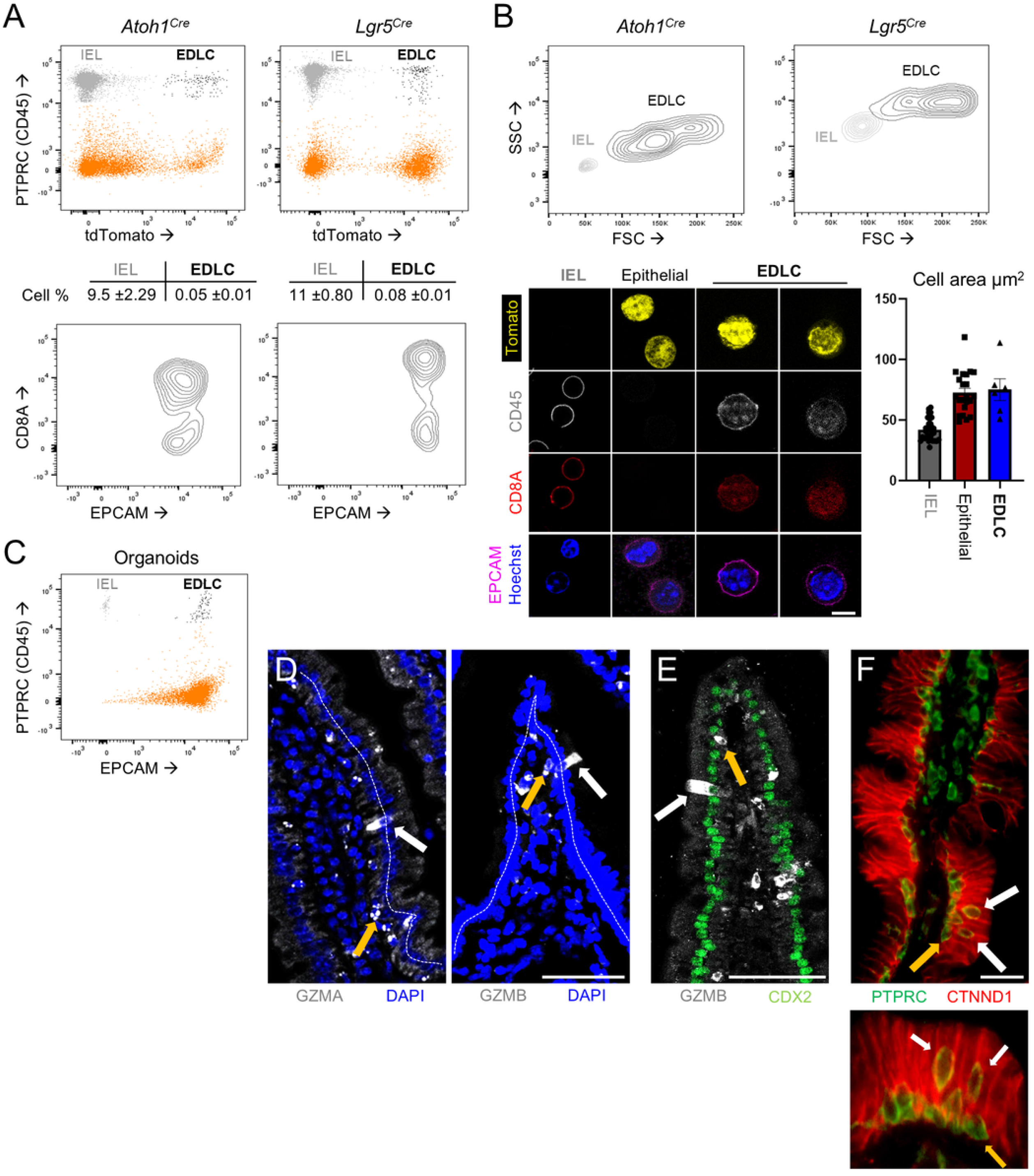
EDLC detection by flow cytometry, with morphology and tissue position distinct from IELs. **(A)** Representative flow cytometry plots for Tom^+^ (*R26*-derived signals after tamoxifen treatment of *Atoh1^Cre-ER(T2)^* or *Lgr5^Cre-ER(T2)^* mice) and PTPRC^+^ (Cy7-conjugated Ab) cells. IELs of peripheral origin (Tom^−^PTPRC^+^ cells, grey) are distinguished from epithelium-derived (Tom^+^ PTPRC^+^ cells, black) EDLCs. Results from 3 (*Atoh1^Cre^*) and 4 (*Lgr5^Cre^*) independent flow cytometry experiments are quantified below. Lower panels show expression of EPCAM (Per-cp-conjugated Ab) and CD8A (Apc-conjugated Ab) in EDLCs gated from the top panel. All EDLCs express surface EPCAM and >60% express surface CD8A. **(B)** Top: representative flow cytometry plots from panel A (n= 7 mice), showing significant differences in forward (FSC) and side (SSC) scatter characteristics between EDLCs and IELs. Bottom: representative images of FACs-sorted cells from *Lgr5^Cre-ER(T2)^;R26Tom* intestines 5 days after Cre activation, showing large Tom^+^ EPCAM^+^ PTPRC^+^ C8A^+^ EDLCs, large Tom^+^ EPCAM^+^ PTPRC^−^ C8A^−^ epithelial cells, and comparatively small Tom^−^ EPCAM^−^ PTPRC^+^ C8A^+^ IELs. Cell areas are quantified in the graph; data represent the mean ±SEM. Scale bar 50 µm. **(C)** Representative flow cytometry plot for EPCAM (Per-CP-conjugated Ab, x-axis) and PTPRC (Cy7-conjugated Ab, y-axis) in cells disaggregated from wild-type mouse intestinal epithelial organoids after 6 days of culture from intestinal crypts. EDLCs accounted for ∼0.12% of all cells and outnumbered the few IELs carried along in the crypt isolates. n= 4 independent experiments. **(D)** Representative micrographs showing GZMA (left), GZMB (right) expression in columnar EDLCs (white arrow), distinguished from abundant IELs at the epithelial base by their larger size and alignment of their nuclei with those of other columnar epithelial cells (denoted by dotted white lines). n= 4 independent experiments, scale bar 50 µm. **(E)** Representative micrographs showing GZMB^+^CDX2^+^ columnar EDLC (white arrow) and GZMB^+^CDX2^−^ IEL (yellow arrow). n= 3 independent experiments, scale bar 50 µm. **(D**) Representative micrograph of human jejunum showing PTPRC^+^ P120CTN^+^ cells (white arrows, EDLCs), distinguished from IELs (yellow arrow) and lamina propria cells by their larger dimensions and nuclear alignment with the columnar epithelium. Boxed area is magnified in the inset below. n= 3 independent experiments, scale bar 50 µm.

If EDLCs are *bona fide* epithelial cells, they should be present in organoids cultured *ex vivo*. Indeed, flow cytometry of small intestine organoids cultured for 6 days in the presence of epidermal growth factor, Noggin, and Rspo1 (ENR medium^28^) identified EpCAM^+^ PTPRC^+^ cells (Fig. 2C). The EDLC fraction, ∼0.12% of all cells, matched the proportions seen *in vivo* and, importantly, far outnumbered the few IELs present in the crypts used to seed organoid cultures. Furthermore, against a backdrop of lamina propria lymphocytes and abundant small IELs packed near the basement membrane, immunostaining detected rare large columnar GZMA^+^ and GZMB^+^ cells present along mouse duodenal villi (Fig. 2D). These Granzyme^+^ cells expressed CDX2, an intestinal epithelium-restricted TF (Fig. 2E), but lacked the common tuft-1 and tuft-2 cell marker DCLK1 (Fig. S2D). Human jejunal epithelium also carried PTPRC^+^ CTNND1^+^ cells, which were distinct from –and larger than– IELs near the basement membrane and more abundant along villi than in crypts (Fig. 2F and Fig. S2E). Mouse and human EDLC nuclei aligned with those of the columnar epithelium, not with basal IELs, further affirming EDLCs as *bona fide* columnar epithelial cells distinct from previously known cells. Neither the mRNA profiles (Fig. 1) nor the appearance of viable EDLCs isolated by FACS or detected in situ is compatible with apoptotic cells.

To characterize EDLCs further, we examined bulk RNA-seq data from *Lgr5^Cre^*-labeled EDLCs and FACS-sorted IELs, both from intestinal crypts (Table S1). Classic IEL markers such as *Gzma/b*, *Ptprc*, and *Cd3e* were expressed at lower levels in EDLCs and, conversely, mRNA levels of α-defensins (which IELs are known to express^29,30^), *Epcam*, and epithelial keratins and TFs were higher than in IELs (Fig. 3A). Importantly, EDLCs lacked transcripts associated with known IEL subtypes, e.g., *Cd4*, *Cd8b1*, *Lag3*, *and Il10ra*, or with tuft cells, e.g., *Dclk1*, *Pou2f3*, *Ptgs1*, and *Plk2* (Fig. 3A). Consistent with known T-cell receptor (TCR) expression in ∼90% of IELs^31,32^ and the presence of B lymphocytes among IELs,^7^ immunoglobulin and TCR transcripts were abundant in IELs; they were, however, notably absent in EDLCs (Fig. 3B). If EDLCs reflect cell fusion, IELs are the most plausible fusion partner and this absence of canonical IEL transcripts, including antigen receptors, further argues against that potential artifact.

**Figure 3.**
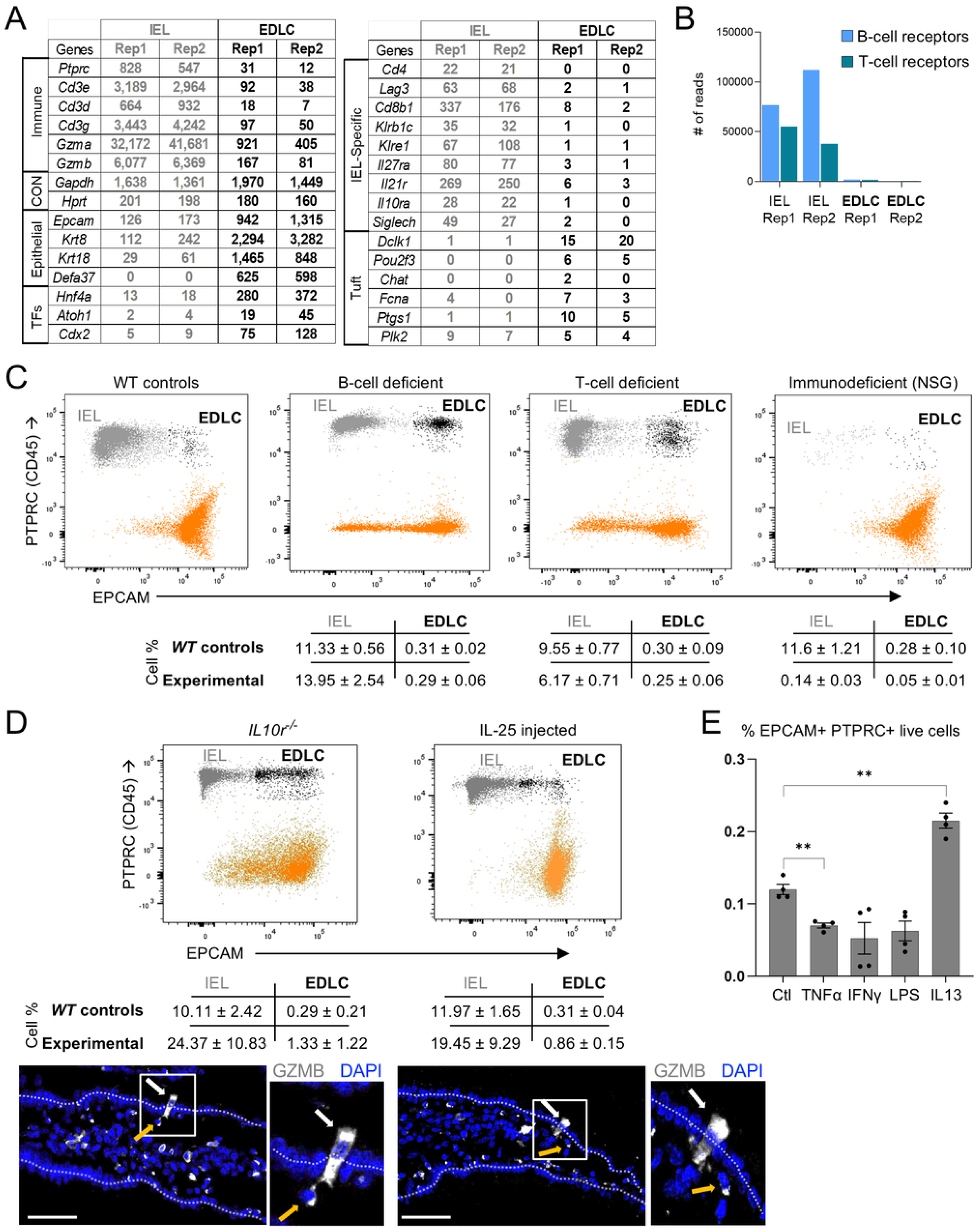
EDLCs differ from IELs in gene expression and their numbers increase in inflammatory conditions *in vivo*. **(A)** Transcript levels (normalized transcripts per million, TPM counts) of selected immune and epithelial genes (left) and IEL-specific and tuft genes (right) from bulk RNA-seq analysis of IELs and EDLCs isolated by flow cytometry from *Lgr5^Cre-ER(T2)^*;*R26Tom* (Fig. 2A and Table S1). n= 2 isolations of each cell type. **(B)** Numbers of immunoglobulin (BCR) and T-cell receptor (TCR) sequence reads from bulk RNA-seq, showing high expression of both in IELs and only background noise in EDLCs. Rep, replicate. **(C)** Representative flow cytometry plots for EPCAM (Per-cp-conjugated Ab, x-axis) and PTPRC (Cy7-conjugated Ab, y-axis) in wild-type (WT), B-cell deficient, T-cell deficient and immunodeficient (NSG) mice. Results from 4 independent flow cytometry experiments for each control-experimental pair are quantified (mean ±SEM). IELs (EPCAM^−^ PTPRC^+^) were moderately upregulated in B-cell deficient mice and reduced ∼35% in T-cell deficient mice, but EDLC (EPCAM^+^ PTPRC^+^) numbers in both conditions were unchanged. In NSG mice, IELs were reduced >80-fold, while EDLCs were reduced <6-fold. **(D)** Representative flow cytometry plots for EPCAM (Per-CP-conjugated Ab, x-axis) and PTPRC (Cy7-conjugated Ab, y-axis) in *Il10r^−/−^* (left) or IL-25 (right) treated mouse intestinal epithelium. Results from 3 independent flow cytometry experiments for each control-experimental pair are quantified. EDLC numbers increased by 4.5-fold in *Il10r^−/−^* and 2.8-fold in IL-25 treated conditions. Bottom: representative micrographs showing GZMB expression in columnar EDLCs (white arrows), distinguished from abundant IELs (yellow arrow) at the epithelial base (n= 3 independent experiments for each state). **(E)** Quantification of the percentage of live EPCAM^+^ PTPRC^+^ cells in cultured intestinal epithelial organoids untreated (Ctl) or treated with tumor necrosis factor α (TNFα, 20ng/mL), interferon y (IFNy, 20 ng/mL), lipopolysaccharide (LPS, 100 ng/mL), or interleukin 13 (IL-13, 15 ng/mL). IL-13 consistently doubled the EDLC fraction, while the other treatments reduced their numbers. Data represent the mean ± SEM. Statistical significance was assessed by one-way analysis of variance followed by Tukey’s test (**, p < 0.01).

To determine if immune cells influence the presence or numbers of EDLCs *in vivo*, we used flow cytometry to examine intestinal epithelium from mice deficient in circulating T or B cells^33,34^ and from *Nod-Scid y* (NSG) mice, which have extreme paucity of all lymphocytes.^35^ IEL numbers (EPCAM^−^ CD45^+^) were modestly perturbed in B-cell^33^ or T-cell^34^ deficient mice, as expected, while EDLC numbers (EPCAM^+^ CD45^+^) were preserved in both cases (Fig. 3C). Whereas IELs were fewer by 2 orders of magnitude in NSG mice, from 11.6% of crypt epithelial cells in wild-type controls to 0.14% of cells, EDLCs were reduced only ∼5-fold (Fig. 3C). This finding further distinguishes EDLCs from IELs but suggests that lymphocyte-mediated host responses might induce or enhance the EDLC phenotype. We tested this possibility in *Il10r^−/−^* mice, which develop spontaneous patchy inflammation,^14,15^ and by injecting wild-type mice with IL-25 daily for 7 days to trigger a type 2 immune response.^13,36^ Average EDLC numbers increased in both cases, by 2.8-to 4.6-fold, and were readily seen by immunostaining (Fig. 3D). We also cultured intestinal organoids in the presence of tumor necrosis factor α (TNFα), interferon y (IFNy), lipopolysaccharide (LPS), or interleukin 13 (IL-13, effector of the type 2 response downstream of IL-25 *in vivo*^13^). Only IL-13 elicited a consistent elevation (doubling) of the EDLC fraction, while other treatments reduced the numbers slightly or left them unchanged (Fig. 3E). Thus, EDLCs increase in response to type 2 immune stimuli and certain inflammatory conditions.

### Chromatin basis of the EDLC phenotype and further exclusion of artifacts

Although these observations together exclude IEL contamination or cell fusion as trivial explanations for EDLCs, other possibilities include atypical IEL phagocytosis and exosome or RNA transfer from IELs to epithelial cells. Because lymphocyte DNA would not stay intact in these scenarios and to provide additional evidence against fused binucleate cells, we analyzed data from assays of transposase-accessible chromatin at single-nucleus resolution (snATAC-seq).^37^ This method stringently excludes nuclear pairs or groups; moreover, profiles of open chromatin could shed light on EDLC origins and the basis for a hybrid phenotype. In snATAC-seq analysis of crypt epithelial cells, *Atoh1*-and *Neurog3*-labeled EDLCs clustered in the same KNN neighborhood (Fig. S3A). This finding affirms their dual phenotype because binucleate cells would separate in the snATAC protocol and cluster with different neighbors. Pseudobulked data identified 3,743 enhancers accessible only in EDLCs, exceeding the numbers specific to EE or tuft cells (Fig. 4A). EDLC-selective sites were largely inaccessible in purified ISCs; conversely, sites enriched in ISCs or goblet/Paneth cells^18^ were weakly accessible in EDLCs, suggesting shared provenance. Importantly, EDLCs shared accessibility at epithelial loci such as *Epcam* and *Krt8* with other cells, but they were uniquely accessible at lymphoid loci such as *Cd8a* (Figs. 4B and S3B). Together, these findings establish epigenetic authenticity and features of EDLCs, implying that certain TFs engage EDLC-specific *cis*-elements to activate lymphoid genes. EDLC-accessible sites were enriched for DNA-binding motifs of IKZF2 and RUNX3 (Fig. S3C). mRNAs encoding these TFs are expressed in EDLCs (Fig. 1D) and their loci are uniquely accessible in EDLCs (Fig. S3D), suggesting that the cellular phenotype might reflect activity of these and other lymphoid TFs.

**Figure 4.**
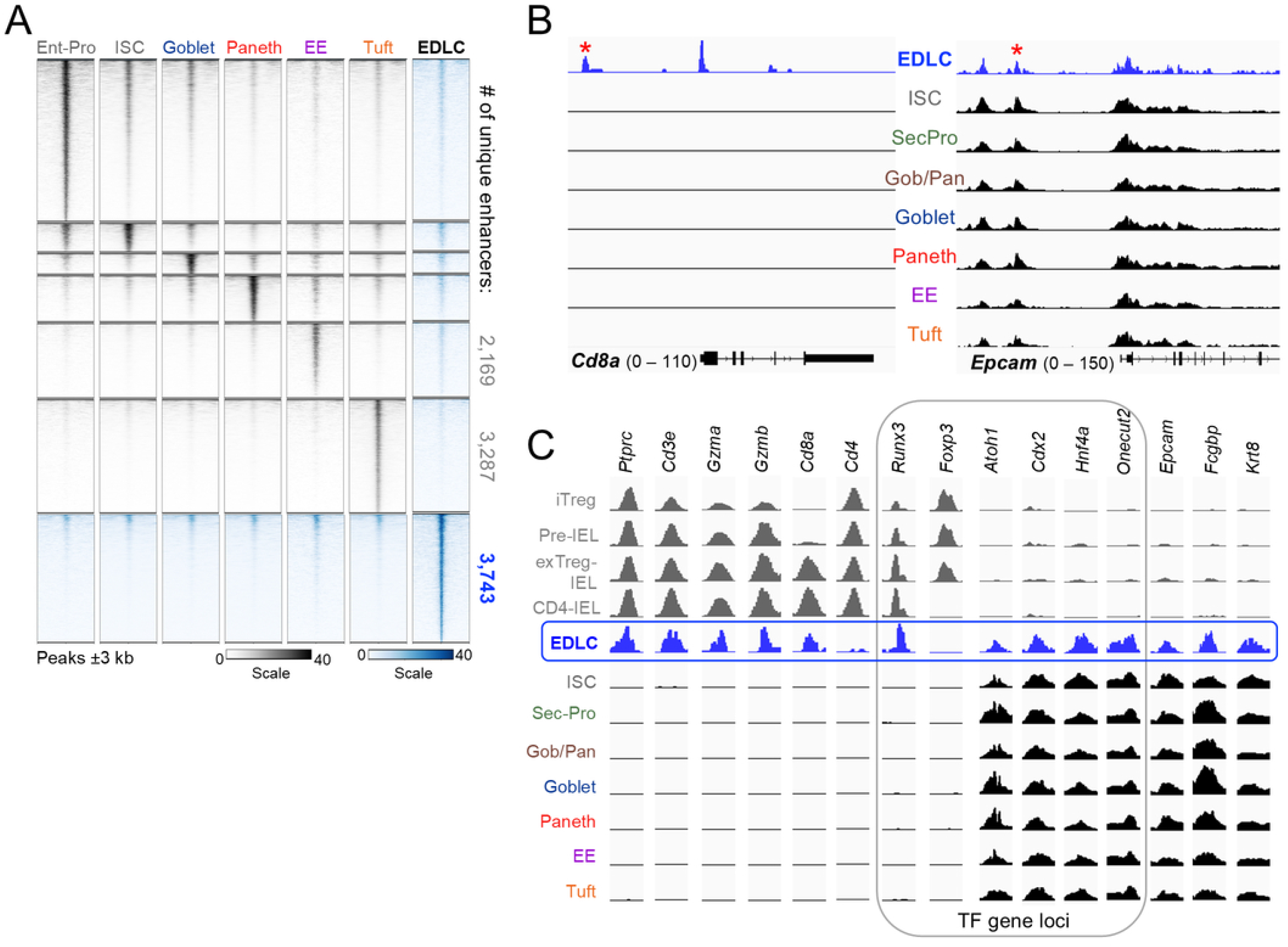
Accessible chromatin at epithelial and lymphoid gene loci in EDLCs. **(A)** Accessible enhancers (>-1 and 2 kb from TSSs) enriched in EDLCs (pseudobulk, Fig. S3A) over other intestinal epithelial cells, extracted from Ref. ^18^ and replicated here in grayscale. EDLCs (blue) carry open chromatin at >3,700 unique enhancers. **(B)** Representative IGV tracks (pseudobulk snATAC-seq) showing chromatin accessibility at *Cd8a* (lymphoid) loci exclusively in EDLCs, while the *Epcam* (epithelial) locus is accessible in all cell types, including EDLCs. Asterisks denote enhancers shown in isolation in Fig. 4C (coordinates listed in Table S2). **(C)** Representative enhancers from snATAC-seq of EDLCs, other *Atoh1*-labeled cells, and bulk ATAC-seq analysis of *Foxp*-labeled *Cd4^+^* IELs (Ref. ^30^) at classic lymphoid, lineage-restricted TF, and epithelial gene loci (chromosomal coordinates listed in Table S2). EDLCs gave robust signals for open chromatin at lymphoid loci other than *Cd4*, which is accessible only in *Foxp*-labeled IELs. Intestinal TF and epithelial and secretory enhancers, accessible in all epithelial derivatives, including EDLCs, are inaccessible in these IELs.

Public snATAC data from IELs are sparse. We compared EDLC data with those from the only IEL populations examined in depth: mature CD4^+^CD8^−^ IELs and their *Foxp3*^+^ iT_reg_ precursors.^30^ Genomic sites uniquely accessible in EDLC chromatin included those in *Cd8a*, *Gzma/b*, *Runx3*, and other lymphoid loci, whereas genes that are not expressed in EDLCs, e.g., *Cd4* or *Foxp3*, were inaccessible (Fig. 4C and Table S2). Conversely, epithelial loci such as *Epcam*, *Fcgbp*, and *Krt8* and intestinal TF loci such as *Cdx2*, *Atoh1*, *Hnf4a*, and *Onecut2* were accessible in EDLCs, but not in *Foxp3*-derived IELs. Of note, sites in *Ptprc* and *Cd8a* were accessible in EDLCs but not in tuft cells (Fig. 4B and Fig. S3B); conversely, tuft-cell loci such as *Dclk1, Pou2f3*, and *Ptgs1*were accessible only in tuft cells and not in EDLCs (Fig. S3D), again affirming non-identity of these cell types. Thus, EDLCs are epigenetically and transcriptionally distinct from any known epithelial cell type, including IELs or tuft cells, and they activate loci classically regarded as restricted to adaptive immune cells.

## Discussion

As the primary barrier against trillions of microbes, the intestinal epithelium provides many layers of protection, including specialized mucus secretion,^38^ antimicrobial products of goblet/Paneth cells,^18^ recruitment of diverse IELs,^6,11^ and antigen-presenting capability in the context of MHC class II antigens.^5,39^ However, only T lymphocytes are thought to express a gene program associated with adaptive immune cytotoxicity. Our findings challenge this long-held view. We identify EDLCs as rare columnar *Ptprc^+^* cells that originate *in situ* from epithelial progenitors, co-express epithelial and lymphoid markers, and are distinct from IELs in numerous respects, including mRNA profiles and lack of antigen receptor transcripts. Large columnar GZMA^+^ and GZMB^+^ cells are distinct from small basilar IELs in immunostained intestinal tissue and in flow cytometry characteristics. Because EDLCs become labeled when Cre recombinase is restricted to various crypt progenitors, they represent either a previously unrecognized cell type within the unitarian model of intestinal cell differentiation^2^ (Fig. 1A) or a cell initially fated for conventional secretory differentiation that subsequently activates lymphoid genes. Our findings do not discriminate between these possibilities.

EDLCs likely evaded recognition previously because the tiny population is vastly outnumbered by IELs. Moreover, to deplete IELs when purifying intestinal epithelium, PTPRC^+^ cells (which mark both EDLCs and IELs) are commonly removed, and our own prior studies of *Neurog3^+^* and *Atoh1^+^* crypt cells^18,20^ reflexively dismissed EDLCs as IEL contaminants. They are, however, reliably detected among rigorously purified *Atoh1-*, *Neurog3*-, or *Lgr5-*labeled cells, not only in single-cell RNA analyses but also by flow cytometry for CD8, PTPRC, and EPCAM; together, these findings exclude the possibility of spurious doublets or phenomena like phagocytosis and efferocytosis.^40^ mRNA release from dead cells (ambient RNA^41^) is an unlikely scRNA-seq artifact because EDLCs express secretory genes but lack a robust transcriptional signature of any specific cell type; moreover, we detected EDLCs by methods independent of scRNA-seq, such as flow cytometry and visualization *in vivo*. Several lines of evidence also argue against cell fusion, a phenomenon well known in tumor and normal, including intestinal epithelial, cells.^26,27,41,42^ The T-cell mRNA program marks T cells as the only plausible fusion partner, yet EDLCs lack antigen receptor transcripts and their numbers are not materially reduced in mice with severe T-cell deficiency. In organoids cultured *de novo* from isolated crypts, where T cells are not present in large numbers for fusion, we detected EDLCs at the same frequency as *in vivo*. Mononuclear EDLCs co-express GZMA and the intestine-specific TF CDX2 *in vivo* and lineage tracing in *Lgr5^Cre^* and *Atoh1^Cre^* mice labeled comparable numbers of EDLCs shortly after reporter activation, a result expected of resident secretory epithelial cells but not from stochastic fusion events. Finally, in snATAC-seq analysis, the nuclei of binucleate cells would separate or single fused nuclei would carry the global chromatin signature of a specific epithelial partner; however, EDLCs revealed a unique hybrid landscape of epithelial and T-cell enhancers. These findings together establish EDLCs as distinct epithelial cells rather than products of stochastic fusion.

Our study does not address EDLC functions *per se*. However, prominent components of the transcriptional program, including *Cd8a*, *Gzma*, *Gzmb*, and other cytotoxic T-cell genes, suggest that they might act as contact-proximal effectors to eliminate certain pathogens or infected or damaged cells. Inflammation and IL-25-induced increase in EDLC numbers *in vivo* is consistent with those functions and suggests that EDLCs may be integral to the type 2 immune response previously characterized in tuft and goblet cells.^13,42^ PTPRC^+^ epithelial cells were not obscure in human jejunal tissue (Fig. 2F and Fig. S2E), were evident in all mice we studied, and relative cell paucity often belies substantial function, as for example in the case of single sentinel cells per colonic crypt.^43^ Therefore, just as innate epithelial immunity counters assorted intestinal microbes^3,4^ and tuft cells combat specific parasites,^44,45^ EDLC features lead us to speculate that they serve in mucosal defense. We propose that certain inflammatory or infectious conditions activate lymphoid TF genes in secretory epithelial cells, that these TFs in turn activate distinct *cis*-regulatory elements to generate partial lymphoid character, and that the resulting dual phenotype provides an additional layer of mucosal protection, including possible cytotoxic capacity. Future delineation of EDLC roles and requirements will add to the growing understanding and harnessing of epithelial-microbe interactions to combat infectious and inflammatory intestinal disorders.

## Materials and Methods

### Animals

*Atoh1^Cre^* mice^46^ were a generous gift from M. Hoshino (National Center of Neurology and Psychiatry, Japan). *R26R^Tom^* (Jax strain 007909) ^25^ *Lgr5^CreER-T2^* (Jax strain 008875) ^1^ T-or B-cell deficient (Jax strains 007850 and 002288) ^33,34^, and *NOD-Scid-y* (*NSG*, Jax strain 005557) ^35^ mice were purchased from Jackson Laboratories. Strains expanded in our colony were maintained on a predominant C57BL/6 background and housed under specific pathogen-free conditions in 12 h light/dark cycles at 23 ±1°C, 55 ±15% humidity, with *ad libitum* access to food and water. Animals were weaned 21 to 28 days after birth and handled according to procedures approved by the Animal Care and Use Committee at the Dana-Farber Cancer Institute. Experimental mice and littermate controls of both sexes were 8 to 28 weeks old at the time of experiments and cell isolations.

### Mouse treatments and intestinal cell isolation

To activate CRE^ER-T2^ in *Atoh1^CreER-T2^*;*R26R^Tom^* or *Lgr5^CreER-T2^*;*R26R^Tom^* mice, we administered 2 doses of tamoxifen, 50 mg/kg (Sigma-Aldrich T5648; 1 mg/mL stocks prepared in cornflower oil) by intraperitoneal (IP) injection 24 h apart. Small intestines harvested 12 h (*Atoh1^Cre^*) or 5 days (*Lgr5^Cre^*) later were rinsed in phosphate-buffered saline (PBS, Gibco, 10010049), then rotated in 5 mM EDTA (Invitrogen, AM9260G) (pH 8) for 45 min at 4°C with manual shaking every 10 min and a change of solution at 30 min. Released epithelium was passed through a 70-µm strainer to separate villi from crypts; each fraction was dissociated into single cells by rotating in 4% TrypLE solution (ThermoFisher A1217702) for 30-45 min at 37°C. Single cells were diluted in DMEM (Corning 17-205-CV) containing 2% FBS. Tom^+^ cells were isolated on a FACSAria II SORP flow cytometer, with elimination of dead (DAPI^+^) cells and pelleted by centrifugation.

### Organoid cultures

Approximately 100 crypts isolated as described above were added to 30 μl Matrigel (Corning, 356231) droplets. The ENR culture medium, described previously ^28^, contained Advanced DMEM (Gibco, 12491015), 1% B27 (Life Technologies, 17504001), 0.5% N2-supplement (Gibco, 17502048), 125 mM N-Acetylcysteine (Sigma Aldrich, A9165), 10% Rspo1-conditioned medium (Harvard Digestive Disease Center), 100 ng/mL Noggin (Peprotech, 120-10C), 100 ng/mL EGF (Peprotech, 315-09-500UG), and 10 μM Y-27632 (LC Laboratories, Y-5301). Organoids were grown for 3 days, then maintained in the same medium (Control) or with addition of tumor necrosis factor α (TNFα, 20 ng/mL, Life Technologies, 315-01A-5UG), interferon y (IFNy, 20 ng/mL, Life Technologies, 315-05-20UG), lipopolysaccharide (LPS, 100 ng/mL, Sigma Aldrich, L4391-1MG), or interleukin 13 (IL-13, 15 ng/mL, Thermo Fisher Scientific, 210132UG) for 3 additional days before harvesting and disaggregation for flow cytometry.

### Flow cytometry and fluorescence-activated cell sorting (FACS)

The gating strategy for analysis of mouse intestinal epithelial cells began with exclusion of debris by plotting forward (FSC) against side (SSC) scatter and selecting the main cell population. Doublets were discriminated by plotting FSC-A (area) versus FSC-H (height) and excluded. DAPI staining was applied to exclude DAPI^+^ dead cells. Cells were stained with antibodies (Ab) against PTPRC (CD45, Invitrogen 25-0451-82, 1:100), CD8A (BD Biosciences 553035, 1:100), or EPCAM (Invitrogen 46-5791-82, 1:100) for 30 min at 4°C in Hank’s Balanced Salt Solution (HBSS, Gibco 14065-056) containing 10 mM HEPES (Gibco, 15630080), 2 mM EDTA, and 0.5% FBS (Corning 35-010-CV). The main gating strategy, with Tom and PTPRC expression, identified Tom^+^ PTPRC^+^ (EDLC) and Tom^−^PTPRC^+^ (IEL) populations; further analysis assessed CD8A expression in the EDLC population (Fig. 2A) and FSC and SSC parameters evaluated cell size (Fig. 2B). Viable (DAPI^−^) PTPRC^+^ mouse EDLCs (Tom^+^) and IELs (Tom^−^) (Figs. 2 and 3) cells were sorted for RNA-seq or microscopic examination using a BD FACSAria II instrument (BD Biosciences, 70 µm nozzle). Data were analyzed in FlowJo software (BD Life Sciences).

### Bulk RNA-sequencing

For bulk RNA-seq, FACS-sorted cells were collected in TRIZOL reagent and RNA was extracted from 5×10^4^ viable (DAPI^−^by flow cytometry) cells using the RNAeasy Plus micro kit (Qiagen, 74030). RNA integrity and concentration were determined using Agilent Bioanalyzer High Sensitivity RNA 6000 pico kits (Agilent, 5067-1513). Libraries were constructed from samples with RNA-integrity number (RIN) >8.5, using SMART-Seq v4 Ultra Low Input RNA kits, and sequenced on the NovaSeq platform (Illumina).

### Immunofluorescence

Mouse intestines were rinsed in PBS, fixed overnight in 4% paraformaldehyde, cycled through a 10%–30% sucrose gradient, and embedded in OCT compound (Tissue-Tek, VWR Scientific 4583); 8-µm sections were prepared using a Leica cryostat. After blocking for 1 h with 0.5% BSA, slides were incubated with FITC-conjugated GZMA (Miltenyi Biotec 130-119-413, 1:100), GZMB (R&D Systems AF1865, 1:200), Alexa Fluor 488-conjugated CDX2 (Abcam ab195007, 1:100), or CD45 (Invitrogen 14-0451-82, 1:200) Ab at 4°C overnight, followed by washes and incubation with anti-goat IgG conjugated with Alexa Fluor 488 (Invitrogen, A-11055, 1:500) or Alexa Fluor 647 (Invitrogen, A-21447, 1:500). After counterstaining with DAPI, slides were mounted with Vectashield (Vector Laboratories, H-1000-10) and imaged on a Zeiss LSM 980 confocal microscope or Leica Thunder Imager microscope. Images were processed using Fiji software.^47^

Human jejunal tissue (Fig. 2) was fixed for 2 days in 10% neutral-buffered formalin, embedded in paraffin, and 5-µm sections were de-paraffinized 3 times in Histo-clear (National diagnostics, HS-200), then rehydrated through a 100% to 75% ethanol series, and treated with target retrieval solution (Dako S1699, pH 6.0) for 15 min in a pressure cooker. After quenching with a blocking solution (Dako, X0909) for 90 min, slides were incubated overnight at 4°C with a mixture of CTNND1 (BD Biosciences, 610133, 1:200) and AF647-conjugated PTPRC/CD45 (Biolegend, 304050, 1:200) Ab in a diluent solution (Dako S3022). After 3 washes in PBS, slides were incubated for 60 min at room temperature with fluorescence-conjugated secondary Ab in diluent solution (Dako S0809), counterstained for 5 min with Hoechst dye (Life Technologies, H3570), washed, and mounted with ProLong™ Gold Antifade Mountant (ThermoFisher, P36930). Immunofluorescence was imaged with a Zeiss Axio Imager 2 using 20X and 40X objectives.

### Data Processing and Quantitative and Statistical Analyses

#### scRNA sequencing

Fastq files were aligned to mouse genome version mm10 and quality control, normalization, and clustering were performed in Seurat package v4.3.0.1 (Ref. ^48^) in R version 4.2.0. Cells with >900 and <3,000 features in duplicate libraries from *Atoh1^Cre^*;*R26^Tom^*(wild-type, *WT*) duodenum, or >600 and <3,000 features in duplicate libraries from *WT* Ileum, and <7% mitochondrial genes were retained. Data on 8,693 and 11,598 ileal cells were merged and integrated using ‘FindIntegrationAnchors’ and ‘IntegrateData’ functions in Seurat (Fig. S1). Cell features were detected with the ‘FindVariableFeatures’ function (2,000 features, selection method “vst”) and unwanted sources of variation, such as mitochondrial gene expression, were regressed out. We used ‘RunPCA’ for principal component analysis (PCA) on the scaled data and reduced dimensionality using Uniform Manifold Approximation and Projection (UMAP) ^49^; *k*-nearest neighbor (KNN) graphs were constructed using the ‘FindNeighbors’ function in Seurat. We used the ‘FindClusters’ function, which implements an optimal Louvain algorithm to group cells together iteratively by nearest-neighbor modules, and the ‘FindAllMarkers’ function with default parameters to identify genes enriched in each cluster, which was assigned a cell identity based on highly enriched genes.

EDLCs from *Neurog3^Cre-ER(T2)^*;*R26^Tom^*intestinal epithelium ^20^ were merged with *Atoh1^Cre^*;*R26^Tom^* cells, with precursor and mature goblet, Paneth, and enteroendocrine cells combined to form single named clusters of each cell type. We used the scDblFinder package, which is particularly effective at detecting heterotypic doublets ^21^, to identify and remove potential doublets (Fig. 1 and Figs. S1 and S3). To address tuft-cell heterogeneity, we sub-clustered tuft cells using the ‘FindSubCluster’ function in Seurat and the top 100 genes expressed differentially in tuft-1 and tuft-2 cells ^22^. Module scores for tuft-2 and EDLCs were calculated using the AddModuleScore function in Seurat.

#### Comparisons with published datasets

We availed of scRNA-seq and snATAC-seq data from *Neurog3^Cre-ER(T2);^R26^Tom^* mouse intestinal epithelium ^20^ (Tom^+^ cells, GEO: GSE183299) and scRNA-seq data from purified IELs (Ref. ^24^, GEO: GSE221006, Fig. 1E) and unfractionated mouse intestinal epithelium (GEO: GSE210724, Fig. 1F). Potential doublets in scRNA datasets were excluded using the scDblFinder package ^21^.

#### Bulk RNA sequencing and TCR and Ig gene inference

Libraries were aligned to the mouse genome version mm10 and transcript counts were generated using the Viper package with default parameters ^50^. Raw counts were normalized as Transcripts Per Million ^51^ (Fig. 3A). T-cell receptor and Ig gene expression (Fig. 3B) were determined using TRUST4 (Ref. ^52^).

#### snATAC sequencing

Fastq files from *Atoh1*-labeled Tom^+^ cells were aligned to mouse genome version mm10. Data were analyzed using the Signac package v1.7.0 in R (https://www.r-project.org/) version 4.1.1. Cells with >2,500 and <30,000 peak fragments, >30% reads within peaks, a genome blacklist ratio <0.075, nucleosome signals <4 and TSS (transcription start site) enrichment score >2 were retained. These filters yielded 15,145 cells from 2 replicates of *WT* duodenum. To capture accurate chromatin profiles from sufficient numbers of EDLCs, 58 EDLCs from *Neurog3^Cre-ER(T2)^*;*R26^Tom^*mice ^20^ (GEO: GSE183299) and 38 EDLCs from *Atoh1^Cre-ER(T2);^R26^Tom^* mice ^18^ (GEO: GSE271615) were combined (Fig. S3A). Data were normalized using the Seurat extension Signac ^53^. Once clusters were annotated, peaks were identified using the ‘CallPeaks’ function in MACS2 (Ref. ^54^) with default parameters, and peaks overlapping with annotated blacklist_v2 regions in mm10 were removed. Enhancers were defined as regions >-1 kb and >2 kb from TSSs, and those present in at least 1% of cells and enriched in any cluster (log_2_ fold-difference >0.25) were identified using the ‘FindAllMarkers’ function in Signac. TF motif enrichment in EDLC enriched peaks (Fig. S3C) was derived using STREME ^55^. Bigwig files were generated and normalized using Haystack v0.5.5 (Ref. ^56^). Heatmaps (Fig. 4A) were produced using the computeMatrix and plotHeatmap functions in DeepTools v3.3.2.0.170 (Ref. ^57^) and read distributions were visualized on the Integrated Genomics Viewer (IGV, v2.5.0, Ref. ^58^).

**Statistical analyses** were performed using GraphPad Prism 9 or R (https://www.r-project.org/). Figure legends specify biological or technical replicates for each experiment. Statistical results are presented as averages, with individual values shown. Error bars represent the SEM.

## Acknowledgments

Supported by NIH grant R01DK082889 (R.A.S.) and generous gifts from the Lind family. J.R.G. is supported by NIH grants R01DK48370, R01DK101332, RC2DK118640, and the Department of Veterans Affairs (I01BX000930). We thank M. Hoshino for the kind gift of *Atoh1^Cre^*mice; S. Madha-Krause for bioinformatic assistance; K. Polyak for helpful discussions and NSG mice; Y.C. Wang and A.J. Lusis for providing Seurat objects from scRNA-seq analysis of IELs; and the staff of flow cytometry and microscopy facilities at the Dana-Farber Cancer Institute. Graphical illustrations were created with BioRender.

## Data Availability

Sequencing data are deposited in series GSE291022 in the Gene Expression Omnibus (GEO) and will be made available to the public before the date of publication. Referees may access the data using the token **udatcyuwprunzix.** We also analyzed publicly available data from GEO series GSE271614, GSE271615, GSE183299, GSE210724, GSE221006, and GSE146292.

Additional information required to re-analyze data reported in this paper is available upon request.

## Author Contributions

S. Bhattacharya, K.B., and R.A.S. conceived and designed the study; S. Bhattacharya, K.B., G.T., O.E., and S.-H.L. conducted experiments; S. Bhattacharya, P.N.P.S., and R.H. performed computational analyses; B.B. and S.S. provided *Il10r^−/−^* mouse intestines; K.H. assisted with fluorescence imaging; J.R.G., S. Beyaz, and R.A.S. supervised various parts of the study; S. Bhattacharya and R.A.S. wrote the paper, with input and approval from all authors.

